# Fish on the platter! Investigating the dietary habits of fishing cats (*Prionailurus viverrinus*) in the Godavari Delta, India

**DOI:** 10.1101/2023.07.29.551074

**Authors:** Giridhar Malla, Paromita Ray, Yellapu Srinivas, Sudhakar Malla, Byragi T. Reddy, Matt Hayward, Kuppusamy Sivakumar

**Author notes:** Corresponding Author: Giridhar Malla. Giridhar Malla < https://orcid.org/0000-0002-2185-2750> Paromita Ray < https://orcid.org/0000-0001-5485-3699> Srinivas Yellapu < https://orcid.org/0000-0002-5412-4717> Sudhakar Malla < https://orcid.org/0000-0002-2259-7499> Matt Hayward < https://orcid.org/0000-0002-5574-1653 > Byragi Reddy < https://orcid.org/0000-0002-8434-0402> Kuppusamy Sivakumar < https://orcid.org/0000-0002-6938-7480>.

## Abstract

The threatened fishing cat (*Prionailurus viverrinus*) is an elusive and medium-sized cat that is adapted to mangroves, swamps, wetlands, and riverine habitats. A close look at the literature indicates that fishing cats are piscivorous, however this is based on very few studies. Understanding the patterns of resource utilisation by species is crucial for assessing their role in ecosystems and in ensuring their conservation. Therefore, our study presents insights into fishing cat feeding patterns from mangroves of the Godavari delta, Andhra Pradesh, India. We collected 303 putative fishing cat scats and conducted analysis using 120 genetically identified scats. Our analysis revealed that fish was the most important prey by fishing cats in the study area, followed by crabs and rodents. The prey composition did not vary significantly between the three seasons but there were differences between the survey years. The niche breadth also varied across the three seasons, from being a generalist in winters to a specialist in summers. Our results suggest that long term conservation and survival of the fishing cats depends on fish populations, which are the main prey of the species and thus recommend the need to protect the fish populations in the Godavari delta and the surrounding riverine habitats. Given the importance of fish to the diet of the fishing cat, the health of waterways throughout their distribution must be one of the focal strategies of conservation action.

## INTRODUCTION

Most of the world’s 41 felid species are elusive and solitary (MacDonald 2010), and are hypercarnivorous (Carbone et al. 1999, Kitchener et al. 2017) – possessing morphological adaptations that enable them to stalk and kill their vertebrate prey with high efficiency. Felids, which act as apex predators in many ecosystems, are often considered as flagship species due to their charismatic status and their dominant roles in shaping the ecosystem functioning (Gittleman 1989, Terborgh 1992, Gittleman et al. 2001, Finke and Denno 2005, Dalerum et al. 2008). However, despite evolving nearly 30 million years ago, the felid group has largely conserved its form and function, except for six species: cheetah *Acinonyx jubatus*– a cursorial hunter; fishing cat *Prionailurus viverrinus* and flat-headed cat *Prionailurus planiceps*– piscivorous; and the margay *Leopardus wiedii*, marbled cat *Pardofelis marmorata*, and clouded leopard *Neofelis nebulosa* – arboreal hunters (Kitchener et al. 2010). Diet plays an important role in determining several life history traits of apex predators (da Fonseca and Robinson 1990, Kok and Nel 2004). To understand predator ecology and prey-predator dynamics in an ecosystem, dietary information is essential (Johnson 1980, Bowland and Perrin 1993, Chock et al. 2022).

Fishing cats are a medium sized (8-17 kg) cat species that along with its close relative the flat-headed cat, are highly adapted to aquatic habitats (Sunquist and Sunquist 2002, Hunter 2019). They inhabit mangroves, swamps, wetlands, highlands, and riverine habitats in parts of South Asia and South-East Asia. Currently, fishing cat populations are threatened because of population declines over the last decade due to the rapid destruction of their habitats. The species is listed as Vulnerable by the IUCN Red List and included in the Schedule I of the Wildlife (Protection) Act, 1972, and Appendix II of CITES (Mukherjee et al. 2016). Due to concerted efforts by scientists and conservationists in the past decade, fishing cats have gathered increased conservation attention, yet limited information is available about their behaviour and ecology from the wild (Nowell and Jackson 1996, Mukherjee et al. 2016).

Fishing cats are solitary and generally hunt their prey alone (Malla and Sivakumar 2014). Literature available on the dietary ecology of fishing cats in the natural habitats is poor in comparison to other cat species (Kitchener 1991, Sanderson and Watson 2011). The few studies that exist have highlighted that they predominantly prey on fishes but also feed on birds, rodents, reptiles, insects, frogs, molluscs, crustaceans, poultry, and carcasses etc. (Mukherjee 1989, Haque and Vijayan 1993, Sunquist and Sunquist 2002, Cutter and Cutter 2010, Malla et al. 2018).

Along the east coast of India, fishing cats occur in mangrove forests and coastal wetlands (Malla 2016). Since mangroves are nurseries and habitats for several fish species, they presumably provide fishing cats with an abundant food supply. Hence, in this study, we aimed to examine the feeding ecology of the fishing cats in the mangroves of Godavari delta, Andhra Pradesh, where a high density of fishing cats has been reported (Malla 2016). These mangroves provide a unique opportunity to study the felid’s diet composition for two reasons. First, this mangrove-lined estuary is a highly dynamic ecosystem with an influence of the daily tidal cycle and the monthly lunar cycle, which drive the movement of fishes within the creeks; the tidal cycle also periodically submerges parts of the mangrove floor especially during the spring high tides. Second, only two other resident carnivore species occur within the study area: golden jackal *Canis aureus* and smooth-coated otters *Lutrogale perspicillata*. Although sporadic reports of jungle cats *Felis chaus* have been reported from the periphery of the mangroves by the local fishers, they rarely use the mangroves.

Giving due consideration to the available literature and the diverse array of potential prey species present in the Godavari mangroves, we hypothesise that the fishing cats in the Godavari mangroves depend on a wide variety of prey species to avoid competition between the co-predators. We specifically aim to gain insights into: 1) resource availability and prey preferences of fishing cats in the study area, 2) importance of individual prey items within the diet of fishing cats, and 3) whether fishing cats tend to be specialists or generalists in a typical mangrove ecosystem. We believe our results will be useful in better understanding the diet pattern and feeding ecology of fishing cats and help in conservation management of the species in mangrove habitats throughout their distribution.

## MATERIALS AND METHODS

### Scat collection

Initially, we conducted a survey along the mangrove forests to identify potential latrine sites or scat deposition sites and confirmed the presence of fishing cats using information collected from local fisherman and camera traps. The study area was surveyed across two study zones of the Godavari mangroves, a) Upper Gowthami-Godavari zone and b) Lower Gowthami-Godavari zone. Owing to logistic challenges, we selected sites located along the creeks of Thulyabhaga (T), Coringa (C), Gaderu (G), Gullarasu (Gu), Ingalayi (In), Savupillarava (Sp), and MettaSinteru (Ms) sub-tidal channels in the two study zones (Figure 1). We collected a total of 303 fresh probable fishing cat scats from the study area between December 2015 and August 2017. Scats were collected within a month across three seasons, viz. summer/dry season (March-May), rainy season/wet (June-October), and winter (December-February). Before each sampling event, we ensured that the older and dried scats were discarded from the latrine sites and hence we collected only fresh scats. Removal of old scats did not affect the activity of fishing cats at the site or from using the site again for scat deposition.

**Figure 1.**
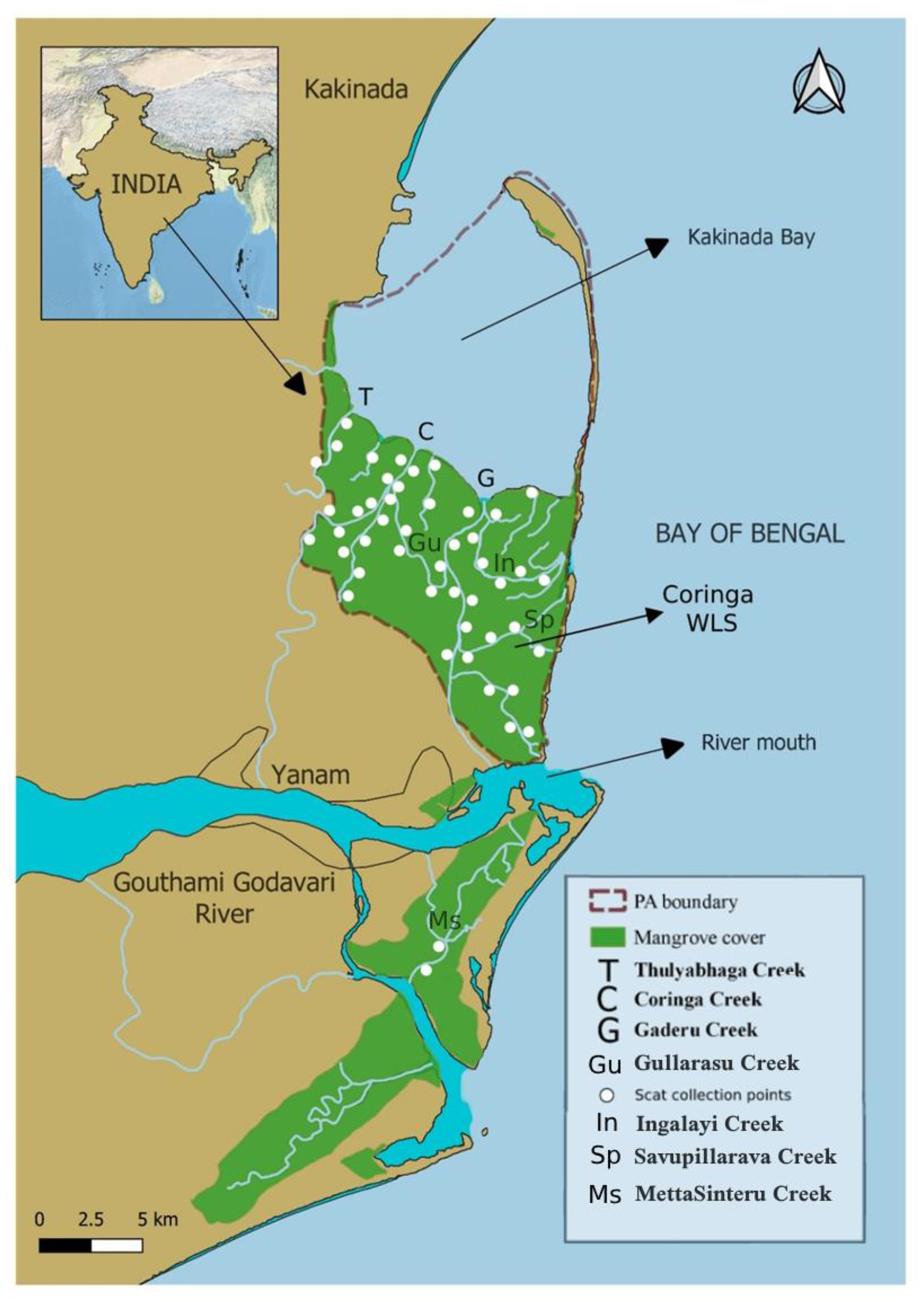
Map showing the scat collection points, along with locations of the creeks, including Thulyabhaga (T), Coringa (C), Gaderu (G), Gullarasu (Gu), Ingalayi (In), Savupillarava (Sp), and MettaSinteru (Ms) subtidal channels in the two study zones.

Fishing cats’ scats were identified based on their shape, size, presence of indirect evidence like pugmarks and scrapmarks, and presence of pungent, fishy smell (Figure 2). Since the study area did not have any other resident carnivore species apart from golden jackal and smooth-coated otters, distinguishing fishing cat scats from the other two species was not difficult, given scats of golden jackal are larger, different in shape and structure (Mukherjee et al. 2004) and smooth-coated otters spraints are generally very smelly, viscous, and sometimes semi-liquid (Prasad 2015). Each scat was collected and stored in a zip-lock plastic bag after labelling the date, creek name, GPS location, and ID number of that particular scat for further genetic and scat analysis. For genetic analysis, we took a small portion of the outer coating of scats and preserved in 90% ethanol solution before shipping to the lab for further storage at –20°C. The remaining portion of the scat was sun-dried for further prey composition analysis.

**Figure 2:**
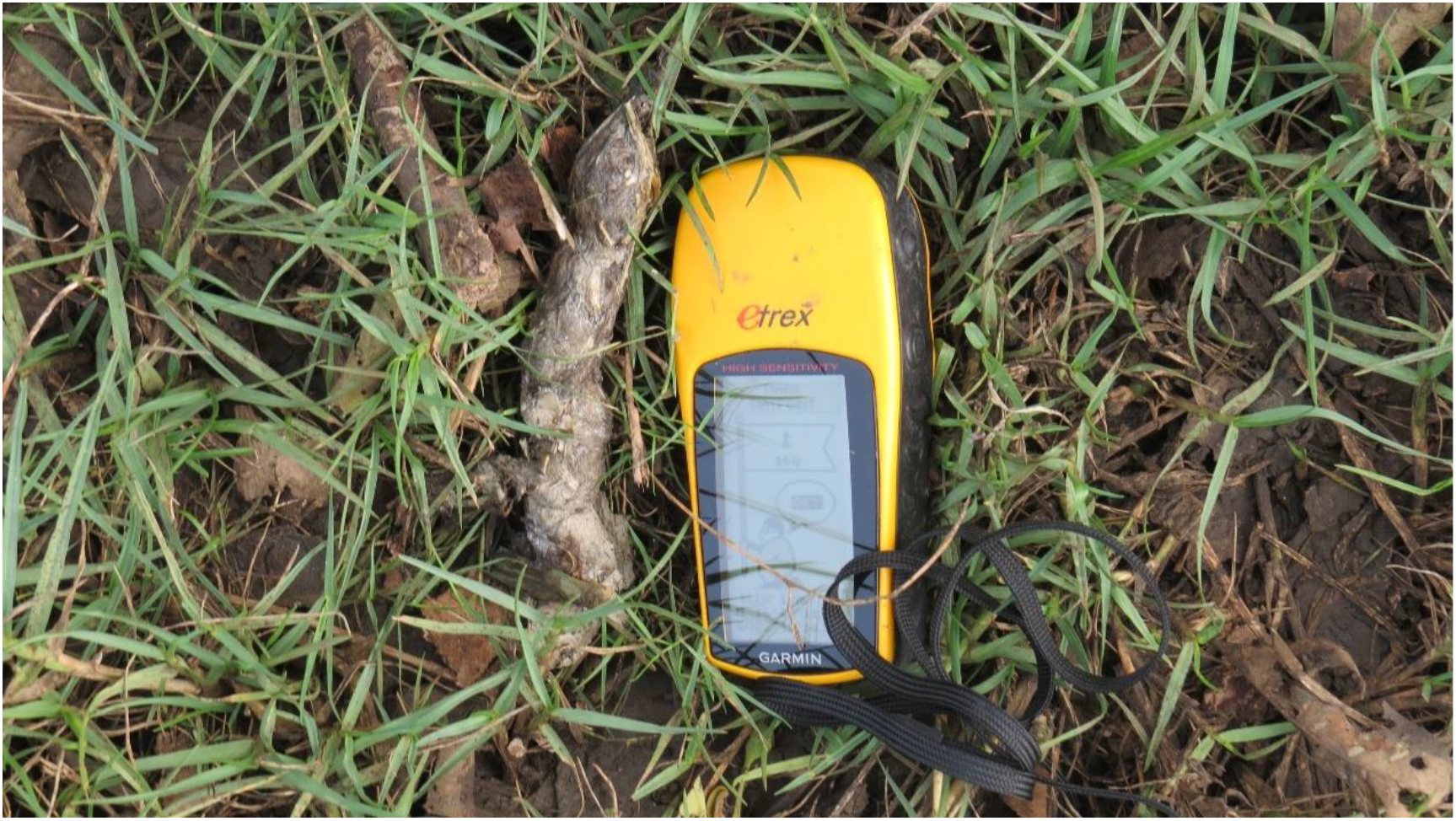
Fishing cat scat collected from the Godavari mangroves, Andhra Pradesh

### Genetic analysis

DNA was extracted using the QIAamp DNA Stool Mini Kit (Qiagen, Hilden, Germany). Extractions were carried out in batches, and negative control was included with every batch to monitor for potential contamination. Species identification of the scat samples was ascertained by using mitochondrial DNA markers: a) carnivore specific Cytochrome b region of 146 bp (Farrell et al. 2000) b) ATP6-ATP8 gene producing 172 bps amplicons (Haag et al. 2009, Trigo et al. 2008). The primers were identified as robust markers for their use of inaccurate species-level identification in the carnivores. Also, the short amplicon size could provide an intact template for amplification especially in case of non-invasive and degraded samples (Chaves et al. 2012). PCR (polymerase chain reaction) amplification for both the primers sets was carried out in a 10 μl reaction volume containing 1X PCR buffer (Thermo Fischer), 2 μg BSA, 4 pM of each primer, 0.5 units of Taq DNA polymerase (Thermo Fisher) and 2-20 ng of genomic DNA. PCR cycling conditions include initial denaturation (5 min at 95°C) followed by 40 cycles of denaturation (the 30s at 95°C), annealing (50s at 55°C), extension (50s at 72°C) and the final extension (10 min at 72°C) in an Eppendorf thermal cycler (Applied Biosystems, Foster City, CA). Necessary measures were taken to avoid contamination and included negative controls during amplification. PCR products were then visualised on 2% agarose gel stained with ethidium bromide. Amplified PCR products were given exo-sap treatment (exonuclease I and shrimp alkaline phosphatase, Thermo Fischer) to remove residual primers before sequencing. The treated PCR products were sequenced bi-directionally using BigDye v 3.1 Cycle Sequencing kit (Applied Biosystems, Foster City, CA). Finally, the sequences generated were validated using GenBank BLAST (http://blast.ncbi.nlm.nih.gov/Blast/) to confirm the species.

### Scat Analysis

Scats were sun-dried, carefully teased apart, and the contents were categorised into mammals, fish, reptiles, insects, crustaceans and vegetation matter. All the non-digested prey remains within the scat including bones, hairs, feathers, exoskeleton of crabs, fish otoliths, fish scales, insect remains, and plant materials were separated and identified to the lowest taxonomic level possible. The prey remains were identified using a 400X microscope and handheld smartphone optics. Fish species were identified to genus level based on the shape and structure of the scales and otoliths based on a references collection obtained from the field and local markets. Since catfishes do not possess scales on their bodies, they were identified using their dorsal and pectoral fin spines and their otoliths. Rodents were identified using hairs, teeth, jaw remains, and tooth alveoli patterns. (Day 1966, Bowland and Bowland 1991). Birds were identified to the order level using their downy barbule structure (Day 1966). Snake species was identified by the scales and the vertebral column bones present in the scats. The crab exoskeletons found in the scats were broken and unidentifiable making it difficult to identify these crustaceans to species level.

### Statistical Analysis

We used the cumulative Brillouin’s diversity index to determine whether the number of scats collected and analysed in this study were sufficient to adequately describe fishing cat diet (Magurran 1988, Glen and Dickman 2006, Hass 2009), using the following equation:

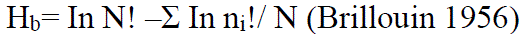

Where H_b_ is the diversity of prey in the scats,

N indicates the total number of prey categories in all samples recorded, n_i_ is the number of individual prey items in the *i*^th^ category

The Brillouin’s Diversity curve was generated by calculating and bootstrapping the samples

10,000 times in two increments. The H_b_ increment curve was then calculated from the incremental change in each mean H_b_ with the addition of two more samples. Finally, the cumulative dietary diversity was plotted against the number of scats analysed to determine if an asymptote was reached or not, and when the incremental change declined to –1%.

Percentage occurrence (PO%) of each prey was calculated by using the number of occurrences of each prey type to the number of occurrences of all prey types multiplied by 100 (Reynolds and Aesbischer 1991, Loveridge and Macdonald 2003). Additionally, presence or absence of each prey species in scats was used to calculate the frequency of occurrence (FO%). The FO% shows the percentage of total samples in which a particular prey item was found, further indicating the importance of different prey types in the fishing cat’s diet (Ackerman et al. 1984, Lessa et al. 2010, Pires et al. 2011). Relative frequency of occurrence was also calculated, which shows the number of individuals of a prey type recorded in the scats relative to the other prey types (Kelly 1991).

Diet breadth and overlap indices were assessed using the major prey categories. Samples were seasonally grouped based on summer, rainy, and winter seasons. Seasonal trophic niche breadth was measured through Levin’s standardised index of food niche breadth (B _standard_) (Erlinge 1981, Henschel 2011, Bobadilla et al. 2022). Niche breadth uses a scale of 0 to 1, representing whether the predator is a generalist (1) or a specialist (0).

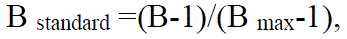

Where: B is Levin’s index (B=1/ p_i_^2^ where p_i_ is the proportion of i^th^ food item identified in every scat); B _max_ is the total number of food categories (Krebs 1999).

Dietary overlap between the seasons was calculated using the Pianka index (Pianka 1973):

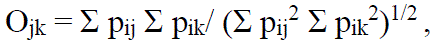

where p is the proportion of the prey category, i for species j and k, values ranging from 0 to 1, indicating total separation/no diet overlap to total/complete overlap.

Additionally, we also looked at the dietary differences among seasons and years on the raw occurrences of prey items using chi-square contingency tables. Fisher’s exact test were used to examine which of the individual prey types significantly differed between seasons. The seasonal diet diversity was calculated using the Shannon–Wiener (H_0_) index and seasonal values were compared using the Hutcheson test (Zar 1996, Silva et al. 2015).

## RESULTS

Out of the total 303 scats collected, we could successfully isolate DNA from 162 scats, of which only 120 scat samples were positively identified as those of fishing cat. The remaining scats were either unidentified or belonged to golden jackals occurring in the study area. Nearly 75% of fishing cat scats (n=90) were collected from specific latrine sites within open spaces (minor to no canopy cover) occupied by below ground vegetation including *Suaeda maritima, Suaeda monoica,* and mangrove seedlings. The remaining scats were found on tree branches, man-made structures like borders of aquaculture ponds, walls adjacent to mangrove patches or human trails. The cumulative dietary diversity curve when plotted against total number of scats showed an asymptote at around 30 samples indicating that our sampling effort was adequate to assess the dietary patterns of fishing cats (Figure 3).

**Figure 3.**
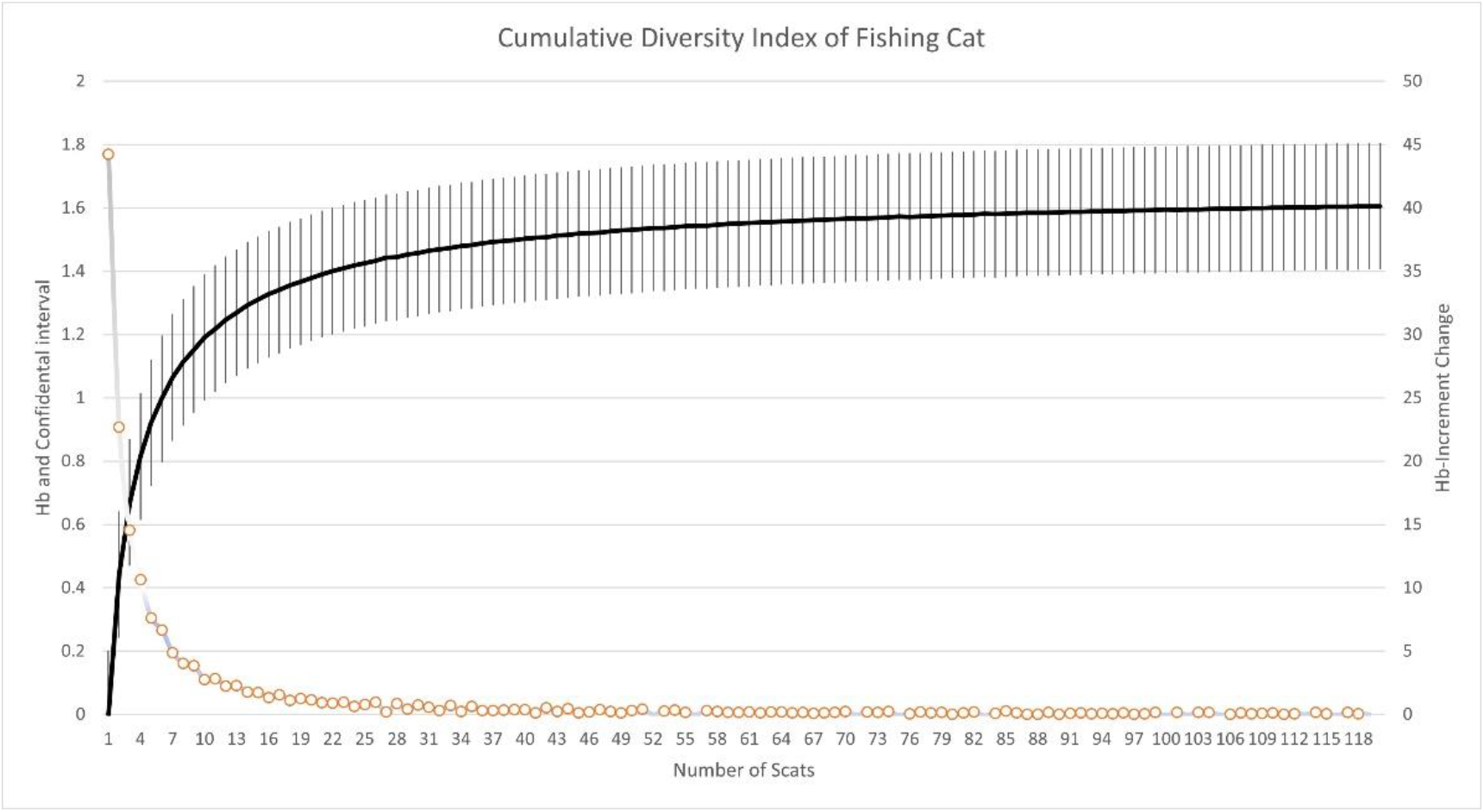
The Brillouin diversity index (Hb) mean and 95% confidence intervals (CI) and incremental change curves. Mean and 95% CI were obtained by resampling with replacement 10,000 times. Mean and incremental change curves reached an asymptote, and the incremental change declined below 1% at ≥ 30 samples.

### Prey composition

We identified 393 different prey items from all the positively identified fishing cat scat samples. The prey items include fish scales, otoliths and bones, feathers and bones of birds, the carapace of crabs, rodent bones, teeth mandible and hair strands, snake bones and scales, and other remains belonging to molluscs and insects. Among the different prey items observed, we identified 16 taxa of plants and animals in the fishing cat’s diet. Out of the 120 scats, 28.3% of scats consisted of only a single prey category, 51.7% consisted of two prey categories, 13.3 % consisted of three prey categories and 0.58% and 0.08% of scats had four and five prey categories respectively (mean = 1.99 ± 0.07 SE). Our analyses revealed that fishes (36.6%) were the most important prey, followed by crabs (17.8%), rodents (16.8%), birds (14.8%), snakes (11.4%) and other including molluscans, insects etc (2.5%).

Fish were the major component of the diet of fishing cats, with consistent numbers observed across all three seasons (n=20 in summer, n=27 in rainy and n=27 in winter). The fish remains observed in the scat samples were identified to seven species with *Mugil cephalus*, *Etroplus suratensis* and *Oreochromis mossambicus* being most important. Fish scales were present in 48 scats, while otoliths and fish bones were present in 34 and 41 fishing cat scats. However, nine scats had all three fish structures: scales, otoliths, and fish bones. Using the otoliths and fin spines, we observed that 12 fishing cat scats consisted of catfishes like *Arius* species and *Mystus gulio*. Crabs occupied second place after the fish category in terms of their importance in the fishing cat diet. Remains of crabs were found in 36 scats, having frequency of occurrence of 30%, of which nine scats had only crab remains. The crab remains could not be identified to species or family levels because in most instances the carapace of the crabs was broken down into smaller pieces. Crabs were recorded more in the rainy season (n=17) and winter (n=15) than in summer (n=4). Rodents occupied third place, occurring in 28% of the fishing cat scat samples, with no major difference seen across seasons (n=5 in summer season, n=13 in rainy, and n= 16 scats in winter seasons).

Birds constituted the fourth most important dietary component and were found in 25% of fishing cat scats with six scats entirely filled by bird remains. Birds were recorded more in winter (n=14) and the rainy season (n=12) followed by summer (n=4). Bird feathers were present in 26 fishing cat scats, bones in eight scats, while six scats contained both bird feathers and bones. Based on the downy barble structures, we identified Indian pond heron *Ardeola grayii*, cattle egret *Bubulcus ibis,* yellow bittern *Ixobrychus sinensis,* white-breasted waterhen*, Amaurornis phoenicurus,* and watercock *Gallicrex cinerea* bird species belonging to Pelecaniformes and Gruiformes. However, 40% of the bird feathers found in the scats remained unidentified due to the poor structure of the remains. Snakes were a minor prey item observed in 23 scats. We observed a major difference across seasons with most identified during winter (n=16), followed by the rainy (n=4) and summer (n=3) seasons. Dog-faced water snake *Cerberus rynchops* was the most commonly consumed snake species.

### Diet composition and Seasonal variation in the diet

Frequency of occurrence of prey species was highest for fishes (61.6 %), followed by crabs and rodents with 30% and 28.4 %, respectively (Table 1). We did not find any significant seasonal differences in either the prey composition or the dietary diversity of fishing cats (Table 2). However, the prey composition did vary between the sampling years (χ2= 82.114, *d. f* = 14, p-value < 0.001). The overall trophic niche breadth for the fishing cats was estimated at 0.56 using the standardized Levins index. Seasonally, the feeding niche of this species ranged from being a generalist in winter (0.75) and the rainy season (0.67) to a fish specialist in summer (0.36). The niche overlap index between summer and the rainy season was 0.92, summer and winter 0.95, and between rainy and winter was 0.89 (Table 3).

**Table 1.** Summary of overall prey items and their percentage of occurrence, frequency of occurrence and relative frequency of occurrence.

| Sl.<br>No | Prey items | Percentage of<br>occurrence<br>(PO%) | Frequency of<br>occurrence<br>(FO%) | Relative<br>frequency of<br>occurrence |
| --- | --- | --- | --- | --- |
| 1 | Fish | 36.6 | 61.6 | 44.4 |
| 2 | Rodents | 16.8 | 28.4 | 13.5 |
| 3 | Reptiles | 11.4 | 19.2 | 6.6 |
| 4 | Birds | 14.8 | 25 | 12 |
| 5 | Crabs | 17.8 | 30 | 17.2 |
| 6 | Others (molluscs and<br>insects) | 2.5 | 4.1 | 6.3 |

**Table 2.** Fishing cat seasonality diet comparisons of dietary composition and diversity using chi-square and Hutcheson test in Godavari mangroves between 2015 and 2017.

|  | DIET COMPOSITION |  |  | DIET DIVERSITY |  |  |
| --- | --- | --- | --- | --- | --- | --- |
| | $\chi^2$ | *d. f* | *P* | *t* | *d. f* | *P* |
| **Summer \* Rainy** | 22.5 | 20 | 0.3012 | 1.479 | 52 | 0.145 |
| **Summer \* Winter** | 28.0 | 25 | 0.3079 | 1.242 | 63 | 0.218 |
| **Rainy \* Winter** | 22.75 | 20 | 0.3012 | 0.3169 | 158 | 0.751 |

**Table 3.** Fishing cat niche breadth and overlap in diet between summer, rainy and winter in Godavari mangroves between 2015 and 2017.

|  | SEASON |  |  |
| --- | --- | --- | --- |
|  | Summer | Rainy | Winter |
| **Niche breadth** | 0.36 | 0.67 | 0.75 |
|  | Summer-Rainy | Rainy-Winter | Summer-Winter |
| **Overlap**<br>**(Pianka's Index)** | 0.92 | 0.89 | 0.95 |

In all three seasons, we observed that fish contributed the most towards the fishing cat’s diet in terms of percentage of occurrence (Table 4). The highest contribution of fishes was recorded in summer (46.5%) further decreased to ∼30% in the rainy season, and ∼25% in the winter season. On the contrary, the proportion of the other three major prey items – rodents, birds, and crabs was the least in summer in comparison to the other two seasons reflecting the increased specialisation that occurs then. The rodents contributed ∼15% to fishing cats’ diet in the rainy and winter seasons and 11.6% in the summer. The overall frequency of occurrence of crabs was higher than the rodents but the frequency of occurrence of crabs (3.3%) was slightly lower in the summer season than rodents (4.2%). The crabs were more frequent in rainy season (14.2 %) than in winter season (12.5%). Birds constituted fourth place in the overall frequency of occurrence. However, in the summer season (3.3%) their frequency of occurrence was the least compared to the winter (11.7%) and rainy (10 %) seasons.

**Table 4.** Seasonal frequency of occurrence (FO) and percent of occurrence (PO) of prey items recorded in Fishing cat scats in the Godavari mangroves between 2015 to 2017. (n=number of occurrence).

|  |  | SUMMER |  |  | RAINY |  |  | WINTER |  |  |
| --- | --- | --- | --- | --- | --- | --- | --- | --- | --- | --- |
| Sl.No | Prey Items | n | FO | PO | n | FO | PO | n | FO | PO |
| 1 | **Mammals** | 5 | 4.2 | 11.6 | 13 | 10.8 | 14.6 | 16 | 13.3 | 15 |
| 2 | **Birds** | 4 | 3.3 | 9.3 | 12 | 10 | 13.5 | 14 | 11.7 | 13.1 |
| 3 | **Reptiles** | 3 | 2.5 | 7.0 | 4 | 3.3 | 4.5 | 16 | 13.3 | 15.0 |
| 4 | **Fishes** | 20 | 16.7 | 46.5 | 27 | 22.5 | 30.3 | 27 | 22.5 | 25.2 |
| 5 | **Crabs** | 4 | 3.3 | 9.3 | 17 | 14.2 | 19.1 | 15 | 12.5 | 14.0 |
| 6 | **Insects** | 1 | 0.8 | 2.3 | 4 | 3.3 | 4.5 | 1 | 0.8 | 0.9 |

## DISCUSSION

Systematic information on diet requirements of fishing cats from the wild is rare. Our study is perhaps among the first to assess the dietary habits of fishing cats in one of the largest river deltas in India. We identified 363 prey items in the analysed scats belonging to 11 taxa, indicating that fishing cats are consuming a diverse and a broad range of prey items in the Godavari mangroves. Our results are in accordance with earlier studies on dietary patterns of fishing cat, showing fishes to be the major component of their diet (Cutter 2015, Haque and Vijayan 1993, Sunquist and Sunquist 2002). While, the remaining prey items including crabs, birds, rodents, and snakes are the next preferred species slightly varied across seasons and years. Our study further highlighted fishing cat to be generalist and an opportunist in the dynamic estuarine ecosystem of Godavari delta and illustrate for the first time how fishing cats increase specialisation at particular times of the year. This increased specialisation may be in response to an increase in fish availability, or it could also be a strategy to reduce dietary overlap with competing predators of more terrestrial prey thereby facilitating coexistence. Interestingly, we also found that the fishing cats preferred defecation of feaces along the aquaculture ponds, local temples, etc., which may be to increase the detectability of the markings while providing a visual amplification (Hayward and Hayward 2010).

### Prey composition

Our study revealed a diverse range of prey items in the fishing cat’s diet encompassing a variety of species, including fish, crabs, rodents, birds, reptiles, molluscs, and insects. Fish constituted the most important prey category with species like *Mugil cephalus* (mullets), *Etroplus suratensis*, and *Oreochromis mossambicus* are preferred among the most common fish species consumed. Mullets are abundant in these mangroves (Sivakumar et al. 2017), which potentially explains their high prevalence in the fishing cats’ diet. Another plausible explanation for the higher dominance of these fish species could be attributed to their feeding habitats. In particularly, mullets are well-known to rely heavily on mangroves for their food and as nursery grounds (Whitfield and Durand 2023). These are benthopelagic fish that are found during low tides, move in large numbers towards shallow water and graze on detritus and algal films covering the exposed mudflats (Cardona 2016). This corresponds well with the unique hunting behaviour of fishing cats we observed during our study wherein they utilized the “sit-and-wait” approach along the exposed mudflats during low tides (Malla and Sivakumar 2014).

Possibly, the fishing cats have adapted to exploit the feeding opportunities presented by the abundance of mullets and estuarine fishes on the exposed mudflats during low tides. Therefore, we believe that fishing cats strategically await the conditions of exposed intertidal zones and optimal water flow to effectively hunt for fishes in the mangrove creeks. This could explain the prevalence of fish species, particularly mullets, in their feaces. Given the fish-rich diversity of mangrove forests, we propose that the ease of catching prey becomes more crucial than prey availability for fishing cats within this dynamic habitat, which accords with the optimal foraging theory. Furthermore, the exposed banks of the mangrove-lined creeks during low tides, especially during extended periods of spring tides, likely provide fishing cats with an advantageous platform for foraging and hunting fish.

Crabs and rodents were the second and third most important prey items after fishes in terms of their percentage of occurrence. The mangroves of Godavari region support over 21 species of crabs (Dev and Bhadra 2008). Since these crabs predominantly forage in the exposed intertidal zones of the mangroves, they are likely an easily accessible prey for fishing cats. During our formal interactions with fishermen and crab collectors in the study area, they reported numerous instances of fishing cats efficiently catching crabs during low-tide periods. Likewise, the mangroves of Godavari are also home to rodents who build nests with leaves on the tree branches. This behaviour of preying on rodents is observed in other small cat species like the jungle cat and the caracal (*Caracal caracal*) (Mukherjee et al. 2004). Although fishing cats primarily feed on fishes, this opportunistic predation on rodents likely helps maintain their basic energetic requirements (Mukherjee et al. 2004).

### Seasonal variation in the diet

Our findings suggest that fishing cats exhibit opportunistic feeding behaviour, targeting a variety of prey items including both vertebrates and invertebrates. Seasonal analysis revealed no significant differences in diet diversity, suggesting that fishing cats diet remain as relatively constant throughout the year. Niche overlap was high between the three seasons, indicating a considerable overlap in the dietary composition of fishing cats during these periods, highlighting the importance of certain prey species.

The overall trophic niche breadth was 0.56, indicating a moderate specialization towards a fish-based diet. On a closer look, the feeding niche of fishing cats varied from being a generalist to a specialist, with the highest food niche breadth observed during the winter season (0.75), followed by the rainy season (0.67), indicating a more diverse diet. In contrast, the summer season displayed the lowest niche breadth (0.36), indicating a relatively more specialized diet dominated by fishes. This pattern was reflected by the increase in contribution of non-fish prey items during the winter and rainy seasons in comparison to the summer season, when fishes constitute ∼46% of total prey. showed that they have enough behavioural plasticity to take advantage of a wide variety of prey species.

The differences observed in the niche breadth can be attributed to the varying conditions of the dynamic habitat throughout the different seasons. During summers, larger area of the intertidal zone along the mangrove creeks is exposed compared to the rainy and winter seasons. In contrast, the rainy season results in large parts of the mangroves being submerged due to the flooded Godavari River, leading to a reduced exposure of the intertidal zone. These changes in intertidal zone exposure likely affect the fishing cats’ hunting capability. A more exposed intertidal surface provides greater access and improved chances to successfully hunt fish, while an inundated intertidal flat limit the access to the prey. Additionally, catching fish may be more challenging for fishing cats during the rainy season due to higher water turbidity caused by the higher sediment load in the flooded waters. Consequently, seasonal variations in the fishing cats’ niche breadth may be influenced by the accessibility of intertidal surfaces and their ability to hunt fish, which is in turn driven by the seasonal changes in intertidal surface area, water turbidity, and other possible environmental factors. These may explain the shift in the fishing cats’ niche breadth from summers, when they specialize on fishes, to the rainy and winter seasons when their diet diversifies and the consumption of non-fish prey increases.

During the winter and rainy seasons, the occurrence of birds in the fishing cat diet was more pronounced compared to the summer season. This variation reflects the seasonal availability and abundance of waterbirds in the study area (Sivakumar et al. 2017). The presence of various coastal habitats, such as mangroves, mudflats, and lagoons, within the study area attract a diverse array of waterbird species, including waders, bitterns, egrets, herons, and waterfowls. Moreover, the Godavari River delta is one of the most important bird areas in India (Islam and Rahmani 2004) and is an important stop-over point for migratory birds along the Central Asian Flyway. This delta annually hosts more than 1% of the global populations of at least 17 waterbird species (Sathiyaselvam and Sreedhar 2015). These bird species exhibit seasonal fluctuations in their abundance, specifically in winters, when they advent over the delta to feed on the intertidal habitats and mudflats. This may consequently affect the relative importance of birds in the fishing cat’s diet across the different seasons.

### Management implications

Our comprehensive three-year study provides invaluable insights into the seasonal diet and feeding behaviour of fishing cats in the Godavari mangroves further adding to the sparse ecological knowledge on this threatened species. The results underscore the dependence of fishing cats on multiple prey resources, with fish being their primary prey. This highlights the urgent need to safeguard fish populations in the Godavari mangroves and the surrounding riverine habitats, with priority given to Mugilidae fishes, which are frequently being consumed by fishing cats. Future studies should investigate the impact of fish resource declines on fishing cat populations as it is likely that their population densities are limited by bottom-up factors like other predators (Hayward et al. 2007). To secure the long-term survival of fishing cats in the Godavari mangroves, it is crucial to establish continuous monitoring initiatives for both fishing cats and fisheries resources. The information we have gathered holds great significance for forest managers as it enables them to ensure an adequate prey base, particularly fish, which is vital for the survival of fishing cats. However, it is important to note that the fishing cat populations within the Coringa Wildlife Sanctuary remain protected, but their numbers outside of this protected area are highly vulnerable to the myriad threats posed to them and their mangrove habitats. While addressing the different challenges, we need a multi-stakeholder and multi-scale approach along with the extensive community-based awareness programs which are equally necessary to educate and engage local communities in the protection of fishing cats and the mangrove habitats. We believe such a collaborative effort will be the first step in fostering long-term survival and conservation of fishing cats in the Godavari mangroves.

## ACKNOWLEDGEMENTS

This study was part of a UNDP – GEF – MoEF &CC – APFD –Wildlife Institute of India project. Giridhar Malla was also supported by MbZ Species Conservation Fund (Project No. 162512790) and WCN Scholarship (2018) for the “*Godavari Fishing Cat Project*”. The authors would like to thank the Director and Dean of the Wildlife Institute of India for providing their support in terms of infrastructure and guidance. The authors would also like to acknowledge the support of the Andhra Pradesh Forest Department, who provided us with the necessary permissions and support to carry out the study. We would personally thank our field guide Prasad Pinapotu, without whom this work wouldn’t have been possible.

